# Local Nearby Bifurcations Lead to Synergies in Critical Slowing Down: the Case of Mushroom Bifurcations

**DOI:** 10.1101/2024.08.08.607203

**Authors:** Mariona Fucho-Rius, Smitha Maretvadakethope, Rubén Pérez-Carrasco, Àlex Haro, Tomás Alarcón, Josep Sardanyés

## Abstract

The behavior of nonlinear systems close to critical transitions has relevant implications in assessing complex systems’ stability, transient properties, and resilience. Transient times become extremely long near phase transitions (or bifurcations) in a phenomenon generically known as critical slowing down, observed in electronic circuits, quantum electrodynamics, ferromagnetic materials, ecosystems, and gene regulatory networks. Typically, these transients follow well-defined universal laws of the form *τ* ∼ |*µ* − *µ*_*c*_| _*β*_, describing how their duration, *τ*, varies as the control parameter, *µ*, approaches its critical value, *µ*_*c*_. For instance, transients’ delays right after a saddle-node (SN) bifurcation, influenced by so-called ghosts, follow *β* = −1/2. Despite intensive research on slowing down phenomena over the past decades for single bifurcations, both local and global, the behavior of transients when several bifurcations are close to each other remains unknown. Here, we study transients close to two SN bifurcations collapsing into a transcritical one. To do so, we analyze a simple nonlinear model of a self-activating gene regulated by an external signal that exhibits a mushroom bifurcation. We also propose and study a normal form for a system with two SN bifurcations merging into a transcritical one. For both systems, we show analytical and numerical evidence of a synergistic increase in transients due to the coupling of the two ghosts and the transcritical slowing down. We also explore the influence of noise on the transients in the gene-regulatory model. We show that intrinsic and extrinsic noise play opposite roles in the slowing down of the transition allowing us to control the timing of the transition without compromising the precision of the timing. This establishes novel molecular strategies to generate genetic timers with transients much larger than the typical timescales of the reactions involved.

## I. INTRODUCTION

Complex systems close to a phase transition or a bifurcation undergo critical slowing down [1, 2], which involves an increase of transient times. This phenomenon lengthens the relaxation times of a system following a disturbance, indicating reduced resilience and increased susceptibility to perturbations. Critical slowing down has been described in a multitude of nonlinear physical systems in optics [3], electronic circuits [4], circuit quantum electrodynamics [5, 6], ferromagnetic materials [7], magnetic quantum phase transitions [8], or models of charge density waves [9]. The investigation of critical slowing down has also been very intensive in the field of ecology, where many works have provided evidence of these phenomena in experimental and field data [1, 10, 11].

The transient time, *τ*, at which orbits reach a small neighborhood of an equilibrium close to a bifurcation typically follows laws of the form *τ* ∼ | *µ* −*µ*_*c*_|^*β*^ for deterministic systems [9, 12–15]. Here, *µ* is the control (bifurcation) parameter, and *µ*_*c*_ is the critical value. The exponents *β* are universal and they are known for local bifurcations. For example, *β* = −1 for pitchfork and transcritical bifurcations [14–16], and *β* = −1/2 for saddle-node (SN) bifurcations [12, 13], with examples of applications in deterministic models of cooperation [13, 17–20]. Similar power laws for transients with exponents −1 have been recently identified for global bifurcations such as the trans-heteroclinic bifurcation introduced in [21, 22], and in bifurcations involving the destruction of quasineutral curves [23]. Recent research has also explored the effects of intrinsic noise on such transients, finding more complicated scaling laws for the SN bifurcation [24, 25]. Such delays for SN bifurcations are caused by the so-called ghosts. This phenomenon appears right after the SN bifurcation where the two equilibria have collapsed and moved onto the complex plane. However, in the vicinity of the region where the SN takes place, the vector field is close to zero and therefore the system undergoes slowing down. Here, we will study a new scenario where the effect of transients of a transcritical bifurcation occurs concomitantly with the ghost phenomena appearing in two close SN bifurcations, which produces a nonlinear (synergistic) increase in transient times.

Phenomena involving the slowing down near SN bifurcations are common in systems and synthetic biology, where cell-type switching dynamics often involve the emergence of bistability and bifurcations [26, 27]. Cellular decisions in biological processes usually require several SNs coexisting within the same system, making it an ideal case study for our research [28, 29]. For instance, during embryonic development, cellular subtypes arise from the cross-repression of genes expressed in each cell type, which together form gene regulatory networks (GRNs). Under spatial signaling, these networks encode the robust spatiotemporal orchestration of cell type specification in the embryo [30–32]. Nevertheless, there is a lack of systematic understanding of the role that different close bifurcations may have in the resulting timing of cell transitions. Therefore, gaining insights into the timing properties of critical phenomena near multiple bifurcations is essential for fully comprehending the temporal dynamics that govern cell decision-making processes.

Slowing down in GRNs has been achieved experimentally in synthetic bistable switches in bacteria [33]. This has opened the door to the proposal of synthetic circuits that perform specific dynamical functions with applications in biomedicine, such as biosensors, customized drug delivery; or the design of responsive materials [34, 35]. Some of these gene regulatory circuits are capable of undergoing several bifurcations while keeping a minimum set of molecular components. This is the case of the AC/DC genetic circuit that exhibits bistability between a stable steady state and oscillations [36]. This multifunctional behavior is paramount in synthetic biology where dynamical instructions need to be encoded with a minimal set of components to avoid the metabolic overload of the cell.

A multifunctional behavior of special interest for synthetic biology applications arises in so-called mushroom bifurcation diagrams, named after its mushroom shape. These diagrams were initially observed in models for neural stem cell differentiation [37, 38], and later in heat-shock protein dynamics in anxiety disorders [39]. The mushroom bifurcation diagram is formed by combining two toggle switches, resulting in four SN bifurcations and three disconnected stable branches [see Fig. 1(b)]. The presence of a mushroom-shaped locus of equilibria provides the system with unique hysteresis properties resulting from two different regions of bistability. Furthermore, it supplies the system with emerging behaviors such as transition-to-Isola states [40], resettable memory, or detec-tion of transient signals [41]. Some of these properties can be related to the potential for isola bifurcation diagrams [42], which have not yet been confirmed experimentally.

**FIG. 1.**
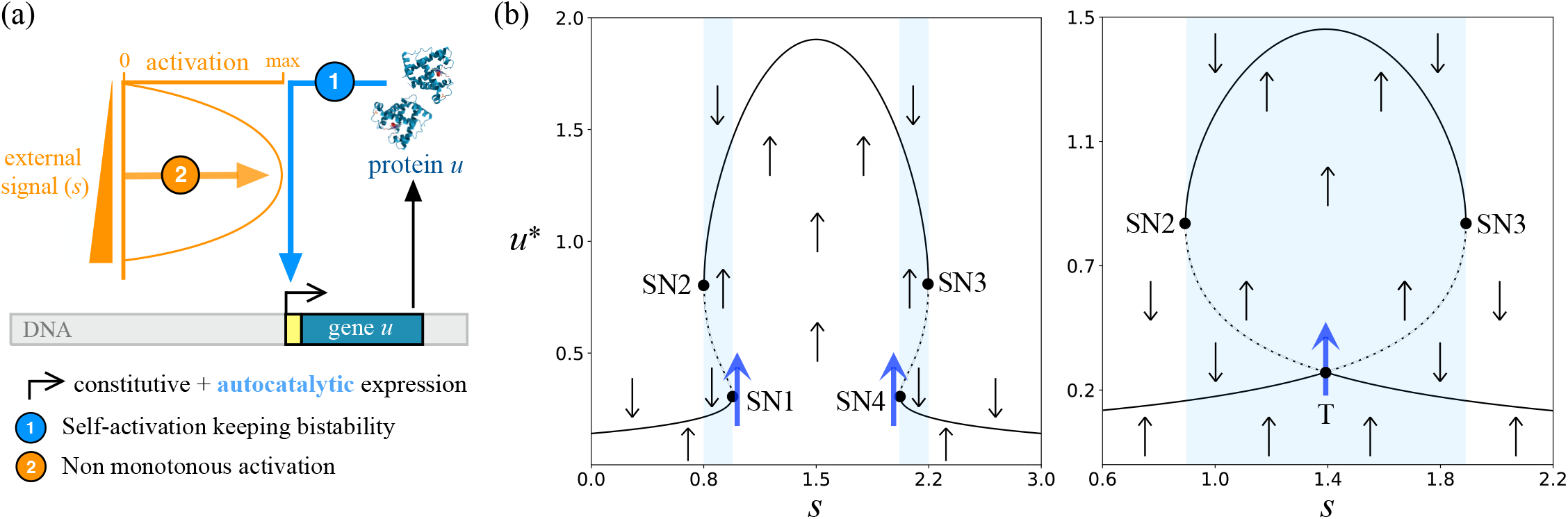
(a) Schematic diagram of the minimal gene regulatory system studied. A gene (*u*) with both constitutive and self-activated expression. The strength of the self-activation of protein *u* is regulated by an external signal (*s*). The effect of *s* is non-monotonic, resulting in maximum autoactivation at intermediate values of *s*. (b) The resulting bifurcation diagram shows the steady-state *u*^*^ as a function of the external signal *s*, corresponding to Eq. (1). The choice of parameters controls the shape of the bifurcation diagram. For high values of *q* the system shows the mushroom bifurcation diagram (left panel with *r* = 0.14 and *q* = 3.0). The neck of the mushroom becomes narrower as *q* is reduced to *q* →*q*_*c*_, resulting in the collapse of SN1 and SN4 towards a transcritical (T) bifurcation (right panel *r* = 0.14 and *q* = *q*_*c*_ ≃ 2.78). The left panel shows the four saddle-node (SN) bifurcations characteristic of the mushroom bifurcation containing two zones of bistability (shaded blue areas with flow direction indicated with grey arrows). The slowing down occurs close to the bifurcations (blue arrows).

Due to this interesting phenomenology, mushroom bifurcations have received special attention lately, mostly in studies trying to dissect the mechanistic rules under which the mushroom bifurcation can emerge in genetic circuits [41, 43–46]. While these studies have focused on the specific shape of the bifurcation diagrams, they only address the steady-state properties of the circuit, without exploring its dynamics. In particular, the possibility of controlling the proximity of two SN bifurcations and exploiting the properties of emerging transients.

To acquire a suitable understanding of the dynamics close to bifurcations in GRNs it is important to consider the effect of noise. Noise may occur intrinsically due to the random nature of biochemical reactions of gene expression, or extrinsically through environmental factors such as noise in cell signaling or cell-cell variability [47, 48]. Intrinsic noise sources occur at different stages of the process of protein expression such as stochasticity in promoter-binding, mRNA translation, or protein degradation; leading to significant molecular heterogeneity [49–51]. Despite its prevalence, molecular noise is usually studied in the context of steady-state dynamics and rarely in the context of transient dynamics. Therefore, further study of the noise on the transient dynamics close to bifurcations is required to fully elucidate the impact of stochastic gene expression on cellular decision-making processes.

Here, we address these questions focusing on the mushroom bifurcation diagram found in a minimal gene regulation circuit. In Section II, we introduce the mathematical model for the mushroom bifurcation diagram, which models a self-activating gene that is regulated non-monotonically by an external signal. In section III we explore the resulting deterministic dynamics of the mushroom and its associated normal form. We present analytical and numerical results for the scaling laws, where, the synergy of two SN bifurcations close to a transcritical bifurcation is introduced. Finally, in section IV, the effect of intrinsic and extrinsic noise in transient times is investigated for coupled bifurcations and compared to single SN bifurcations. Section V is devoted to conclusions.

## II. 1-D MUSHROOM MODEL

### A. Deterministic mean-field description

To investigate the properties of the mushroom bifurcation diagram, we propose a minimal nonlinear one-dimensional system that exhibits this behavior, i.e. a model of a self-activating gene, *u*, which is controlled by an external signal, *s*, [Fig. 1(a)]. This external signal has an activating non-monotonic effect, inducing maximum auto-activation of *u* at intermediate signals *s*. In the deterministic limit, this system is described by the ordinary differential equation:

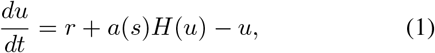

where *u* has a constant basal/leaky expression, *r*, and a linear decay (with a rate set to one without loss of generality). The self-activation term, *H*(*u*), given by the positive Hill function

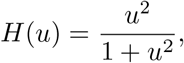

is regulated by the external function *a*(*s*) = *sq*−*s*^2^. To ensure bistability *r* must satisfy

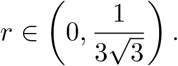

For 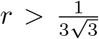, the system is monostable, rendering it unsuitable for our study. The parameter *q* controls the shape of the signal function *a*(*s*), which remains positive (*a*(*s*) > 0) in the interval [0, *q*]. The quadratic shape of *a*(*s*) captures the incoherent nature of the signal, which induces a maximum activation of *u* at an intermediate signal *s* = *q/*2, and fully represses *u* self-activation at the extreme values *a*(0) = *a*(*q*) = 0 [see Fig.1(a)]. The shape of the resulting bifurcation diagram can be controlled with parameter *q*. At high enough values of *q* the system exhibits a mushroom bifurcation diagram with two zones of bistability (Fig. 1(b) left). As *q* is reduced at a critical value *q*_*c*_, two of the SN bifurcations collide undergoing a transcritical bifurcation, and after this collision, the mushroom shape transforms into an isola which becomes disconnected from the lower stable branch (Fig. 1(b) right, and Fig. 9).

### B. Stochastic description

Intrinsic noise is studied by explicitly implementing the reactions of protein production and degradation controlling the dynamics of the absolute number of proteins, *U* = Ω*u*, where the system volume, Ω, captures the relevance of the fluctuations. The rate functions in table I describe the stochastic model. These rates result in a birth-death process of protein production and degradation with propensities *W*_1_(*U*) and *W*_2_(*U*), respectively. In the deterministic limit, where noise is negligible (Ω → ∞), Eq. (1) is recovered.

**TABLE I.** Intrinsic noise model. Protein *U* is produced and degraded with the propensities *W*_1_ and *W*_2_, respectively.

| Elementary process | Propensities | Stoichiometry | Description |
| --- | --- | --- | --- |
| $\emptyset \longrightarrow U$ | $W_1 = \Omega \left( r + \frac{a(s)U^2}{(\Omega^2 + U^2)} \right)$ | +1 | regulated protein production |
| $U \longrightarrow \emptyset$ | $W_2 = U$ | -1 | linear protein degradation |

We additionally incorporate the effect of extrinsic noise as stochastic fluctuations around a given input signal *s*_0_. To do so we describe the stochastic dynamics of signal *s* as an unregulated process with a constant production rate (*R*) and linear degradation rate (*R/s*_0_). Defining the total number of signaling molecules as *S* = *s*Ψ, the signal fluctuations result from another birth-death process where the volume parameter Ψ controls the intensity of the extrinsic noise (see table II). The deterministic limit for the extrinsic noise is also recovered in the limit Ψ → ∞ where the signal remains constant (*s* = *s*_0_).

**TABLE II.**
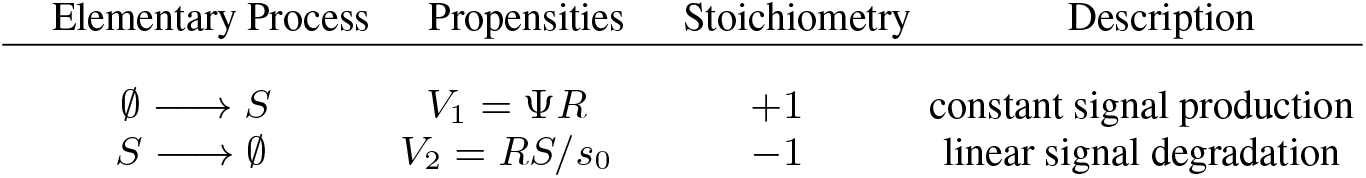
Extrinsic noise reactions. Both reactions with propensities *V*_1_ and *V*_2_ introduce fluctuations around the state *S* = Ψ*s*_0_. Simulations along this manuscript use *R* = 1.

## III. DETERMINISTIC SLOWING-DOWN OF MUSHROOM BIFURCATION DIAGRAMS

In this section, we investigate the deterministic dynamics arising in the mushroom bifurcation diagram focusing on the case where the lower SN bifurcations, SN1 and SN4, approach each other Fig. 1(b). To do so, we first introduce a normal form displaying two SN bifurcations able to collapse into a transcritical one. This model will be useful to obtain analytical results that will allow us to analyze the behavior of the full genetic model described by Eq. (1). The numerical results for this section have been obtained using a 7^*th*^ − 8^*th*^ order Runge-Kutta-Fehlberg-Simó method with automatic time step size control and local error tolerance 10^−15^.

### A. Normal form

For completeness, we start recalling that the normal form of a single SN bifurcation is given by 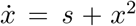. In this study, we will consider the normal form

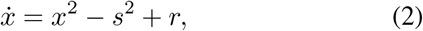

depending on a control parameter *s* and a structural parameter *r*. This normal form describes a dynamical system that can exhibit two SN bifurcations that can collide into a transcritical bifurcation (see Fig. 2). For *r* > 0, the system has two separate SN bifurcations. As *r* decreases, these SN bifurcations approach each other leading to a transcritical bifurcation at *r* = 0, when the SN bifurcations merge [see Fig. 2(a)]. Finally, for *r* < 0, there are no bifurcations, as illustrated in Fig. 2(b).

**FIG. 2.**
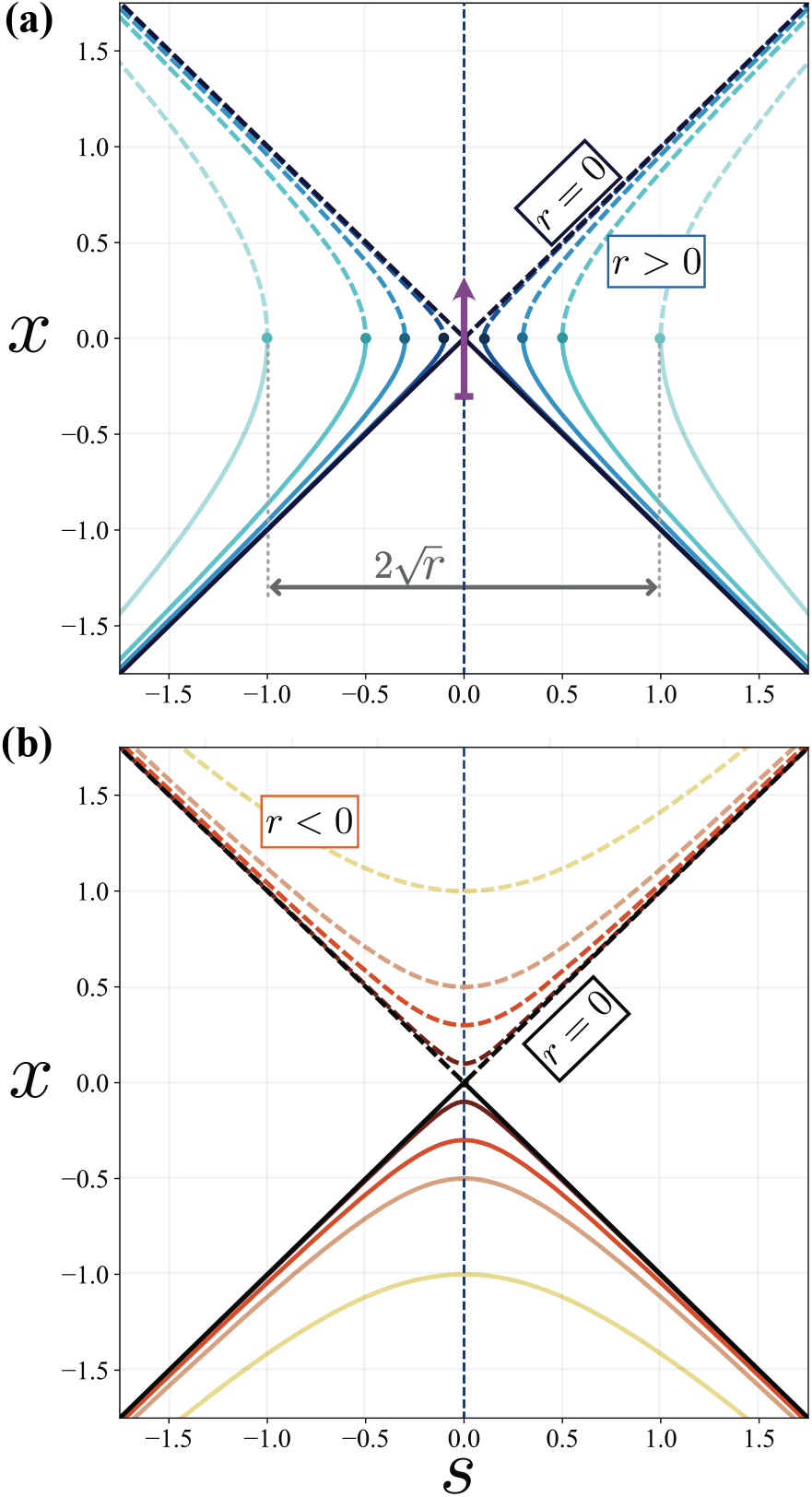
(a) Bifurcation diagram obtained from the normal form Eq. (2). The solid and dashed lines correspond to stable and unstable steady-state branches, respectively. (a) As *r* decreases (for *r* > 0) two saddle-nodes (SN, circles) at a distance 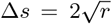 move towards each other colliding in a transcritical bifurcation at *r* = 0 (diagonal lines). For all values of *r* > 0 the system can evolve from *x* < 0 to *x* > 0 (purple arrow) with a transient time that will depend on the location of the SN bifurcations. (b) For *r* < 0 no bifurcations are found.

Differentiating the right-hand side of Eq. (2) reveals that equilibria of the system are stable when they are located at *x* < 0, and unstable for *x* > 0. In particular, for the case *r* = 0, Eq. (2) simplifies to 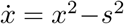, containing the steady-state branches *x*^*^ = − *s* and *x*^*^ = *s*. Thus, both branches exchange stability at the origin, indicating that the system undergoes a transcritical bifurcation.

To study the role that both SN bifurcations have on the transient times, we will consider the transition from *x* = −*ϵ* to *x* = *ϵ* for a fixed value of *r* > 0, being *ϵ* an arbitrarily small real value. The scaling law of the SN bifurcation of the normal form model [12, 13] is examined by fixing a value of 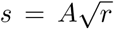, leading to an analysis of the extended transients close to each SN point, i.e. 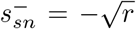 (left SN point) and 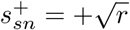 (right SN point). Here, *A* ranges from −1 to 1 re-sulting in values of *s* that span the full range of values between both SN bifurcations. The duration *τ* of the transients as the parameter 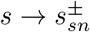 (see Fig. 2), is computed using *s*^2^ = *A*^2^*r*, from the integral

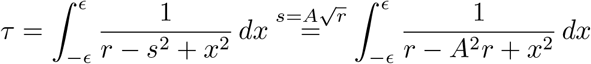

Obtaining

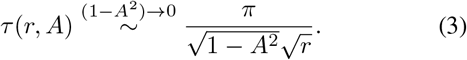

Notably, in contrast to the normal form for a single SN bifurcation [52], notice that the times (3) have a pre-factor dependent on *r*, specifically 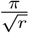, which indicates that as the two SN bifurcations converge i.e., *r* → 0^+^, the slowing down is significantly amplified, thereby lengthening the transient times *τ* while keeping the inverse square root law.

To assess the slowing down close to the transcritical bifurcation, we study a trajectory along the symmetry axis *s* = 0 or equivalently, the neck, between the two SN bifurcations, while progressively approaching the SNs to their convergence, equivalently, as *r* → 0^+^, as illustrated in Fig. 2(a). The integral representing the time scale of transient times *τ* (*r, s*) is expressed as follows:

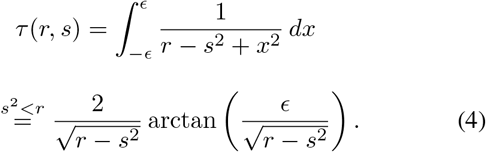

Finally, we obtain

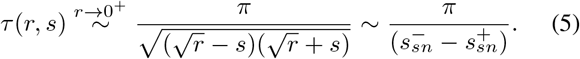

Note that we recover the scaling law for the transcritical bifurcation: 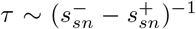. It is also noteworthy that one could follow a similar strategy to study the transients at the formation of the isolas [see Fig. 9(d,e)] [41, 43]). To analyze this case, one should use the normal form:

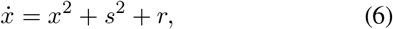

where the value of *r* controls the appearance of the isolas (for more details on a criterion to identify isolas and transcritical bifurcations, see Appendix A).

### B. Gene regulatory system

We now study the dynamics of the minimal genetic system introduced in Eq. (1) [see Fig. 1(a)], regarding transients and their scaling properties.

Without loss of generality, we will focus on transient times close to SN1. To study how delays are affected by nearby bifurcations, we will study the change in transient times as SN1 and SN4 approach each other. To do so, we introduce the quantities 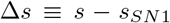, and 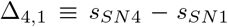, where *s*_*SN*1_ and *s*_*SN*4_ denote the values of the signal *s* at which the bifurcations SN1 and SN4 occur respectively [see Fig. 3(a)]. The magnitude Δ*s* allows us to analyze the effect of a given signal relative to the value at which SN1 occurs. This is the basis for examining the scaling law in a single SN bifurcation and understanding how other bifurcations affect the transient times. On the other hand, Δ_4,1_ controls the proximity of SN1-SN4, allowing us to study synergistic effects between both SNs and the transcritical bifurcation occurring when Δ_4,1_ approaches 0. As Δ_4,1_ → 0^+^ we can derive the power law with exponent − 1, associated with the transcritical bifurcation. This behavior is affected by SN1 and SN4, by introducing pre-factors that extend their respective timescales (see Appendix B). The neck width Δ_4,1_ can be controlled by keeping *r* constant and decreasing parameter *q*, changing the signaling effect on the autoactivation.

**FIG. 3.**
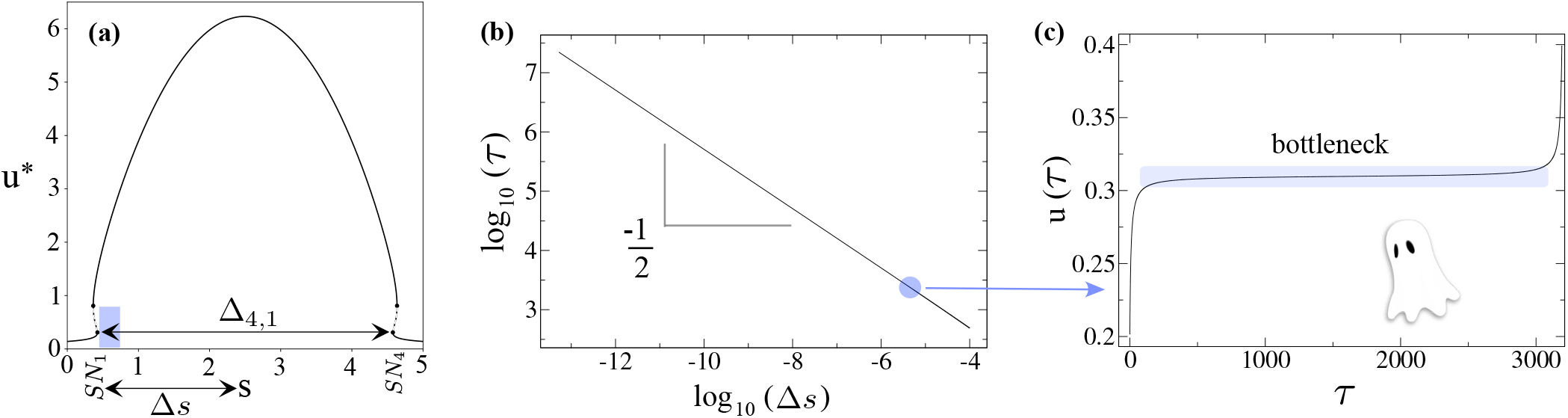
(a) Mushroom bifurcation diagram showing the steady-state *u*^*^ as a function of the signal *s* for the self-activating gene model given by Eq. (1) when the lower saddle-nodes SN1 and SN4 are far from each other (*q* = 5, *r* = 0.14) corresponding to a mushroom neck distance Δ_4,1_ ≃ 4. (b) Transient times *τ* as a function of the distance signal Δ*s* across the slowing-down re close to SN1 (corresponding to the shaded zone in panel (a)). Transient times are calculated computationally as the time *τ* required to reach *u*(*τ*) = 0.4 from an initial state *u*(0) = 0.2 (SN1 is located at *u*^*^ = 0.31). (c) Characteristic bottleneck of ghost transients for log_10_(Δ*s*) = −5.6 [blue circle in panel (b)].

As expected, when both SN bifurcations are far from each other (Δ_4,1_ > 1), numerical analysis of the transient time shows the well-known inverse square-root power-law scaling for transient times (*τ*∼ Δ*s*^−1/2^) [see Fig. 3(b)]. That is, just after the bifurcation, the orbits experience delays and long plateaus associated with delayed transitions [Fig. 3(c)] [9, 12, 13, 30]. For an analytical study of the SN bifurcation scaling law in the self-activating gene model, see Appendix C.

We posit that the ghosts corresponding to SN1 and SN4, which are influenced by the slowing down associated with the transcritical bifurcation, will have a non-linear influence on each other so that transient times will increase synergistically.

To validate this hypothesis, we examined the slowing down at signal values close to SN1 (Δ*s* ≪ 1) for scenarios in which the neck of the mushroom is very narrow [Δ_4,1_ ≪ 1, see Fig. 4(a)]. The resulting slowing down is much longer than what we would expect from a simple additive effect of two SNs. This synergistic amplification can be seen in Fig. 4(b,c) where transients are displayed for different mushroom necks (Δ_4,1_). By comparing the resulting transients with the expected values from simply adding the transient times of two independent SN (blue line in Fig. 4(b,c), and Fig. B.1), we observe that the resulting transients can be orders of magnitude longer. In Section B in the Appendix we compare the delays for single local bifurcations and consider additivity in transient times with the multiple bifurcations described above.

**FIG. 4.**
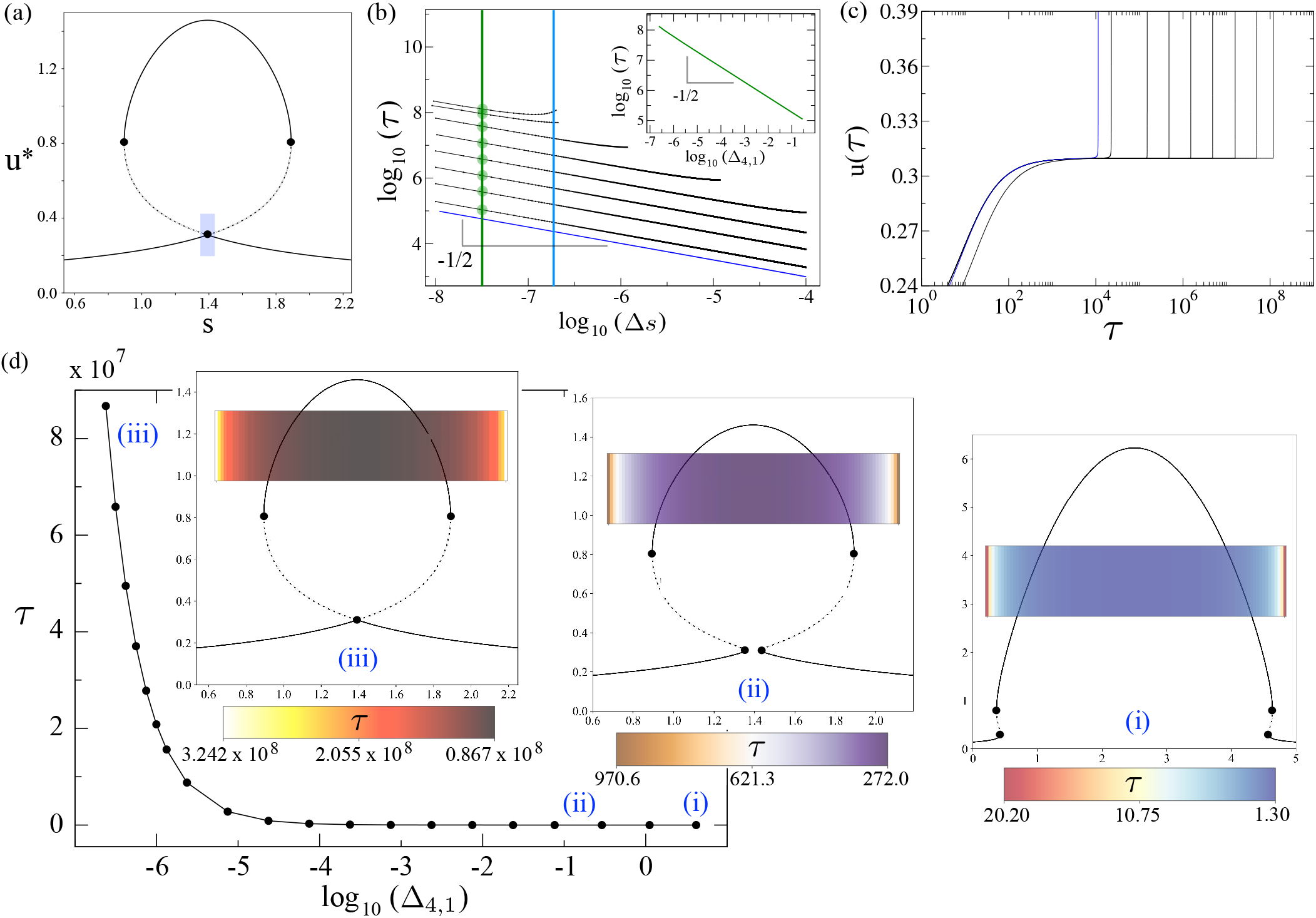
(a) Mushroom bifurcation diagram computed from Eq. (1) with a very narrow neck using log_10_(Δ_4,1_) ≈ − 6.62. (b) Transient times as a function of Δ*s* for different neck sizes (from bottom to up) log_10_(Δ_4,1_) ≈ {0.62, −0.54, −1.63, −2.63, −3.63, − 4.63, −5.63, −6.34, −6.62}, other parameters and conditions are the same as in Fig. 3. The results show a -1/2 scaling law with transient times longer than what would be expected by the additive effect of two independent SNs (blue line, obtained by doubling the time expected by a single SN in the case of a wide-neck mushroom). (Inset) The same scaling law is obtained for the transient times as a function of neck width for a fixed value of log_10_(Δ_*s*_) = − 7.5 (vertical green line and dots). (c) Time series for the same neck sizes and conditions as in panel (b) for a fixed signal value log_10_(Δ*s*) = − 6.7, corresponding to the vertical blue line in panel (b). (d) Transient times keeping Δ*s* = Δ_4,1_/2 shown in log-linear scales. Three examples are displayed: (i) *q* = 5; (ii) *q* = 2.786; and (iii) *q* = *q*_c_ + 10^−14^, corresponding to log_10_ (Δ_4,1_) = 0.62, log_10_ (Δ_4,1_) = −1.12, and log_10_ (Δ_4,1_) = −6.62, respectively. The color maps indicate the duration of transients in the gap between SN1 and SN4, which is computed by dividing this gap into 10^2^ equal intervals.

Furthermore, the smaller the gap (smaller values of Δ_4,1_), the slower the transition, deviating further from additivity. This reveals a synergistic amplification of transients. On the other hand, the scaling law *τ* ∼ Δ*s*^−1/2^ is preserved, suggesting that the slower transient results in an increase in the prefactor of the scaling law that cannot be explained merely by the appearance of the transcritical bifurcation but requires the proximity of both SNs.

In an attempt to clarify the behavior of the prefactor and understand better the synergistic effects, we analyzed the prefactor in the scaling law. We did so by studying the increase of the transients as the width of the neck varies for a fixed value of Δ*s* [green dots and inset in Fig. 4(b)]. This analysis showed that the prefactor also follows an inverse square-root power-law behavior. Such behavior is consistent with the results of the normal form [Eq. (2)] found in the previous sec-tion. Furthermore, it can be derived analytically for the mushroom model of Eq. (C6) (see Appendix C).

To further study the effect of the coupling between both SN bifurcations as they approach each other, we studied the slowing down for signals exactly at the middle point between both saddle nodes Δ*s* = Δ_4,1_/2 for different values of Δ_4,1_ [Fig. 4(d)]. Similar results were found, where for a narrower mushroom neck (log_10_(Δ_4,1_) ≈ −6.6), transient times are seven orders of magnitude slower compared to a wider neck. This observation reiterates the significant influence of the interaction between the two collapsing SN bifurcations. Figure 4(d) shows (in log-linear scale) how the delays increase as the width of the mushroom neck closes. This curve follows the scaling behavior with a power exponent −1 associated with the transcritical bifurcation (see the case with black data in Fig. B.1 for the same results in the log-log scale). This indicates that the slowing down effect on the system is influenced not only by the two ghosts but also by the transcritical bifurcation. An alternative approach to derive this result is to express *τ* as a coupling of the SN slowing down as a function of *a* in Eq. (1), along with the decreasing slope of *a*(*s*). Specifically, we can write *τ* ∼ (*a*(*s*) −*a*_*c*_(*s*))^−1/2^ ∼ (*s* − *s*_*c*_)^−1^, using the fact that near the transcritical bifurcation, *a*′(*s*) = 0 and *a*″(*s*) < 0.

## IV. STOCHASTIC DYNAMICS IN MUSHROOM BIFURCATIONS

To investigate the effect of noise on transients in the self-activating gene model we will initially consider separately the effects of intrinsic (demographic) and extrinsic (environmental) noise following the reaction schemes developed in tables I and II. Then, we will study the system considering both sources of noise simultaneously.

### A. Intrinsic noise

Intrinsic noise results in fluctuations of *u*, which can induce switching in the bistable zones of the mushroom [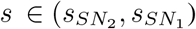 and 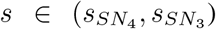. See blue areas in Fig. 1(b)]. Thus, intrinsic noise leads to an expansion of the signal values over which gene expression levels can transition to the upper branch of the mushroom (starting with the initial condition *u* = 0). This effect is observed in any genetic bistable switch [30, 54]. In our particular case, this implies that our study for transient times needs to expand to signals beyond the regions studied in the deterministic scenario. Figure 5(a) shows the transition times calculated as first passage times towards the upper branch of the mushroom. For all values of signal and noise intensities, intrinsic noise accelerates the transient concerning the deterministic scenario. This effect is more dramatic close to the SN bifurcations where the deterministic times diverge while there is still a finite stochastic transient time. For larger noise intensities (lower Ω) the transients are shorter. This leads to an almost constant transition time along the whole mushroom neck for high intrinsic noise (Ω = 10). This behavior is in line with the fact that intrinsic noise allows the system to overcome the ghost, escaping the regime for which 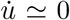. Studying the coefficient of variation (the standard deviation of the mean first passage time divided by its mean), we observe that the precision of the distributions decreases with noise. This suggests that intrinsic noise can be detrimental to encoding precise transient times.

**FIG. 5.**
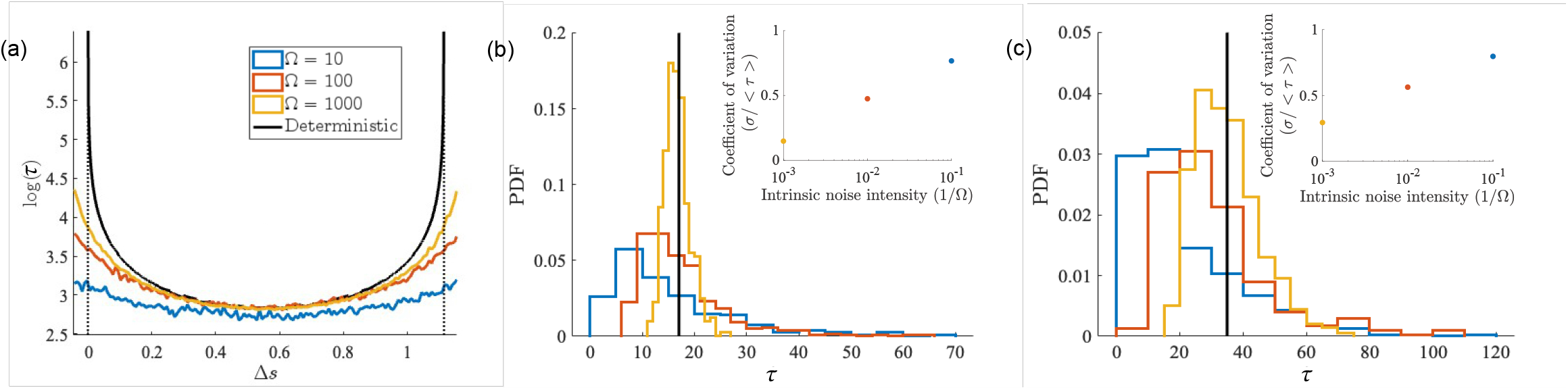
Transition times, *τ*, computed as the first passage times towards the upper branch of the mushroom *u*_*τ*_ = 1.4, starting with initial condition *u*_0_ = 0. Deterministic transients (black solid line) are compared with stochastic transients for different noise intensities: Ω = 10 (blue), Ω = 10^2^ (red), and Ω = 10^3^ (yellow). Parameters are *r* = 0.14, *q* = 3, resulting in a mushroom neck Δ_4,1_ = 1.15. (a) Transient times as a function of the signal. Positions of saddle-nodes SN1 and SN4 (Δ*s* = 0 and Δ_4,1_ = 1.1), are indicated by dotted lines. (b) Histogram showing the transition times probability distributions for a fixed value of signal Δ*s* = Δ_4,1_/2. (c) Histogram of transition times for a smaller mushroom neck Δ_4,1_ = 0.6 (*q* = 2.85) at a signal Δ_*s*_ = Δ_4,1_/2. Simulations are performed using Gillespie algorithm [53] using the reaction schemes described in tables I and II.

As in the deterministic case, constricting the neck of the mushroom results in longer transients that are not erased by intrinsic noise, even in scenarios with high intrinsic noise [Fig. 5(c)]. Similarly, the trend of decreasing precision of transient times with noise intensity is also preserved.

### B. Extrinsic noise

Alternatively, we can study the role of extrinsic noise by considering noise in the signal, while keeping the deterministic dynamics of the gene regulation (no intrinsic noise). Interestingly, for signal regimes between the SNs, extrinsic noise significantly slows down the transient compared to the deterministic scenario [see Fig. 6(a)], in opposition to what was observed for intrinsic noise. This stochastic slowing down increases with increasing extrinsic noise intensity (lower values of Ψ). However, as in the intrinsic noise scenario, extrinsic noise also expands the signal region over which gene expression can transit toward the upper branch of the mushroom. Consequently, close to the SNs where the deterministic transient time diverges, extrinsic noise accelerates the transient.

**FIG. 6.**
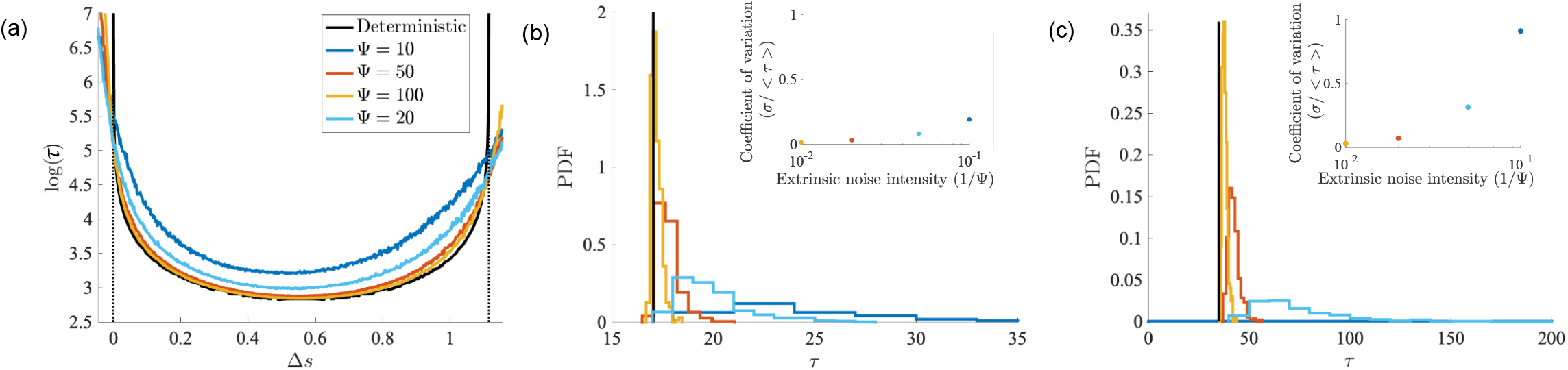
Transition times, *τ*, for the mushroom under the same parameters and conditions as in Fig. 5. Deterministic transients (black solid line) are compared with stochastic transients for different extrinsic noise intensities: Ψ = 10 (blue), Ψ = 20 (cyan), Ψ = 50 (red), and Ψ = 100 (yellow). (a) Transition times at different Δ*s*. (b,c) Histograms showing probability distributions of transient times for two mushroom neck sizes Δ_4,1_ = 1.15 and Δ_4,1_ = 0.6.

Analysis of the distribution of transient times under extrinsic noise shows a similar effect to intrinsic noise in which higher noise is translated into a higher coefficient of variation. Nevertheless, for a large range of noise intensities the coefficient of variation remains very low (below 0.1), suggesting that extrinsic noise does not have a high impact on the variability of the transients [see Fig. 6(b)]. High coefficients of variation (C.V.>0.5) are only observed in limit cases with narrow mushroom necks and high extrinsic noise [see Fig. 6(c)].

### C. Full stochastic model

Given the opposite effects of intrinsic and extrinsic noise within the neck of the mushroom Δ_4,1_, we sought to investigate the behavior of the system when both sources of noise are introduced simultaneously, a situation expected to happen in a real experimental setup. To do so, we analyzed the transient times for different combinations of intrinsic and extrinsic noise (Fig.7). The results show that mean transient times can be faster or slower than the deterministic time, depending on the combination of intrinsic and extrinsic noise intensities. Notably, one can keep the same timing as the deterministic scenario by balancing both sources of noise over a wide range of values of noise intensities [Fig.7(b)]. However, this relationship is different for the coefficient of variation of the timings, where the precision is primarily controlled by the intrinsic noise intensity [Fig.7(c)]. This suggests a mechanism by which one can achieve a specific transient timing with a prescribed precision by balancing the intrinsic and extrinsic noise intensities.

**FIG. 7.**
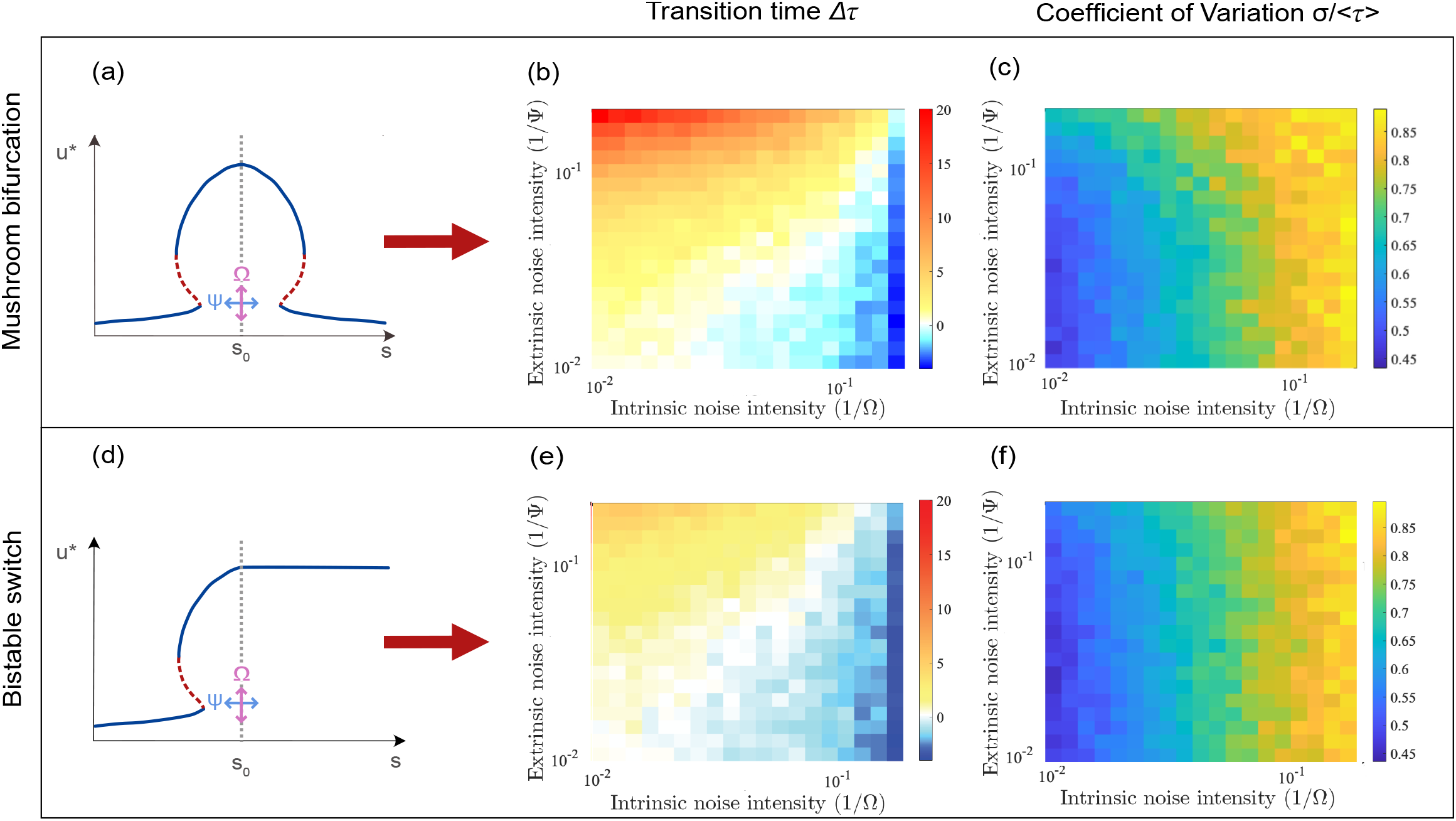
Transition times and variability measures for mushroom bifurcations (a–c) and bistable switches (d–f), for *r* = 0.14 and *q* = 3 with initial expression *u*_0_ = 0 to threshold *u*_*τ*_ = 1.4 for variations in extrinsic stochastic noise with Ψ and variations in intrinsic noise Ω at Δ_4,1_/2 = 0.5575. Note that for the deterministic system, the transition time is *τ*_*det*_ = 17.0439. Panels (a) and (d) show schematic diagrams for the mushroom bifurcation and the bistable switch, respectively. The dotted grey line at *s*_0_ corresponds to Δ_4,1_/2, and the pink and blue arrows correspond to the effective direction of intrinsic and extrinsic noise, respectively, as controlled by Ω and Ψ. (b) and (e): Difference in mean transition times from deterministic time Δ*τ* = ⟨*τ*⟩ −*τ*_*det*_. (c) and (f): Coefficient of variation *σ/* ⟨*τ*⟩. Simulations are performed using Gillespie algorithm [53] employing the reaction schemes described in tables I and II.

Some of the timing dynamics for the noisy system for the mushroom bifurcation can be attributed to the existence of SN bifurcations independently of the mushroom bifurcation diagram. To understand the coupling effect of the SNs characteristic of the mushroom we compared the transient average time and precision with the case in which a single SN is present (Fig. 7). Specifically, we studied a bifurcation diagram corresponding to a traditional bistable switch built by keeping the mushroom bifurcation diagram up to its symmetry point Δ*s* = Δ_4,1_/2 and fixing a constant steady state for higher values of the signal (see Appendix D for details). Before considering the case with mixed noises we can study the effect that intrinsic and extrinsic noise have independently on the transient times of the bistable switch. As expected, intrinsic noise transient is unaffected for values of Δ*s* < Δ_4,1_/2, where the transients of the mushroom and bistable switch are identical [Fig. 8(a)]. On the other hand, intrinsic noise transients are independent of the signal for Δ*s* > Δ_4,1_/2 [Fig. 8(a)]. These results confirm that intrinsic noise effects only depend on the steady states available at a particular value Δ*s*, independently of the bifurcation diagram. This is not true for extrinsic noise, where mushroom and bistable switch transients are different for all values of Δ*s*, including values Δ*s* < Δ_4,1_/2 for which both bifurcation diagrams are identical [Fig. 8(b)]. This is the result of extrinsic noise allowing the system to sample different signal values, and thereby affected by localized slowing down effects near the SNs. Since the effect of the SNs is to slow the transients, extrinsic noise is slowing both the bistable switch and the mushroom bifurcation. This effect is stronger for the mushroom where two SNs contribute to the slowing down dynamics [Fig. 8(b)]. As the width of the neck increases, it is expected that the difference in transition times between both diagrams will decrease, as a signal will have a lower probability of attaining values close to the SNs.

**FIG. 8.**
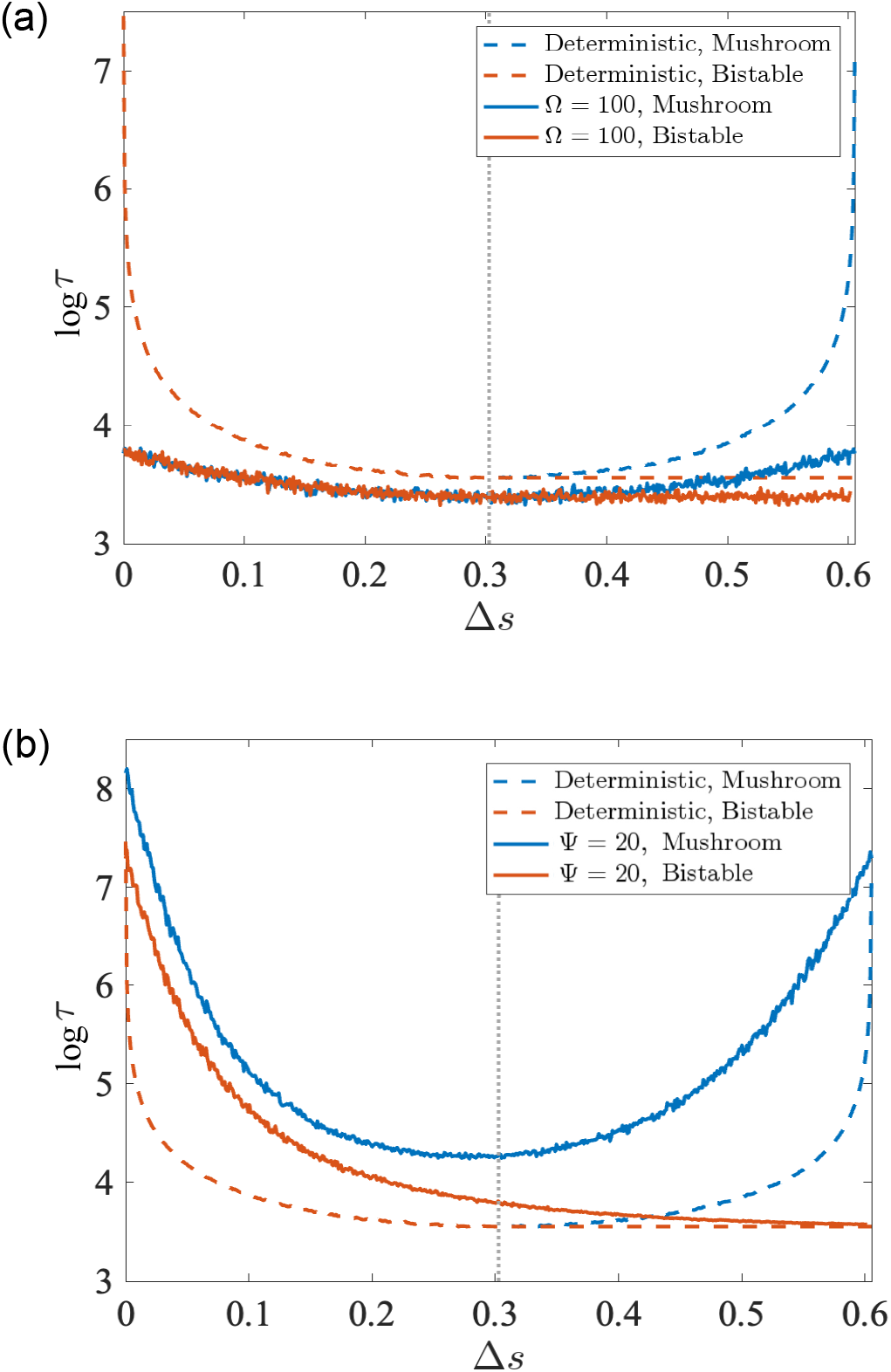
Transient times at different signal values for the mushroom bifurcation (blue lines) and the bistable switch (red lines) with *r* = 0.14, *q* = 2.85 i.e., Δ_4,1_ = 0.6. The symmetry axis of the mushroom where the bistable switch and mushroom start to differ is located at Δ*s* = Δ_4,1_/2 = 0.3 (dotted grey line). Transients of the corresponding simulations in the deterministic case are also plotted for reference (dashed lines). (a) Comparison of deterministic and stochastic transition times for a system with intrinsic noise. (b) Comparison of deterministic and stochastic transition times for a system with extrinsic noise.

Next, we aim to understand whether balancing intrinsic and extrinsic noise to control the mean transient time and precision is a specific property of the mushroom configuration as shown above, or if it can also be achieved in the bistable switch. Comparison of the transition times for different noise intensity combinations (Figs. 7d–f) show that at Δ_4,1_/2, the transient times in the bistable switch diverge from the deterministic case with a similar trend to the one observed in the mushroom. However, the change in transition time is much larger for the mushroom bifurcation. In particular, the mushroom allows for a more dramatic slowing down of the transition, achieving transient timescales that can be twice slower than the deterministic time. Meanwhile, when comparing the coefficient of variation between the mushroom and the bistable switch, we find that the coefficient of variation does not change significantly across the models [cf. Figs. 7(c,f)]. All in all this suggests that the mushroom bifurcation allows for a better control of the slowing down dynamics in stochastic scenarios.

## V. CONCLUSIONS

Transients are known to become extremely long close to critical transitions in a phenomenon known as critical slowing down. This phenomenon has implications in physical and biological systems introducing dynamics much slower than the timescale of the rates composing the system [1, 3–8, 10, 11]. The transient times *τ* near deterministic bifurcations typically follow universal scaling laws of the form *τ* ∼|*µ* −*µ*_*c*_|^*β*^ [9, 12]. Such exponents have been identified in both local [13–15] and global [23] bifurcations. Despite critical slowing down and transients’ scaling phenomena have been widely investigated over the past decades for single bifurcations, either local or global, how dynamical systems behave close to nearby bifurcations remains unexplored.

In this article, we have studied critical slowing down in systems exhibiting mushroom bifurcation diagrams. This bifurcation diagram is usually found in mathematical models of gene regulation [41, 43–46] and, for appropriate parameter values, has four saddle-node (SN) bifurcations. The simple model presented considers a gene that self-activates itself and is regulated by an external signal. By tuning biochemical parameters it is possible to arbitrarily control the distance of the two SN bifurcations forming the neck of the mushroom. By further reducing this distance, the neck of the mushroom eventually closes until both SN bifurcations collapse into a trans-critical one. For this system, we show that the slowing down found right after a SN bifurcation largely increases when the two SN bifurcations approach each other. Moreover, the inverse square-root scaling law remains preserved, as shown for SN bifurcations studied with delay differential equations [18]. We also provide analytical and numerical evidence of this synergistic effect by examining a normal form with the same phenomenology i.e., two SN bifurcations merge into a transcritical one.

In the context of gene regulatory networks (GRNs), our findings provide a mechanism to control the timing of an event, achieving timescales slower than those of protein production and degradation. This suggests a novel approach to constructing genetic timers. Traditionally, GRN transients are overlooked in systems biology, with a primary focus on steady states. However, the timing of transitions is crucial for understanding GRN function. For example, differences in embryonic developmental timing between species often arise from the same reaction networks with nearly identical rates, highlighting the importance of understanding time control in de-velopmental processes [55].

One of the defining features of GRNs is the significant impact of noise due to the low number of molecules involved. Our study demonstrates that specific bifurcation diagrams, which can be achieved with minimal GRN topologies, not only exploit these transients but also buffer the effects of noise. Importantly, we show that intrinsic and extrinsic noise have opposite effects on timing. This discovery enables the achievement of specific timings with prescribed precision by controlling intrinsic noise (e.g., through plasmid number regulation in synthetic circuits) and extrinsic noise (e.g., by modulating signal noise).

Furthermore, while our study explores the one-dimensional case, it opens the door for future research to investigate scenarios involving multiple genes. Examining how different bifurcations interact could unveil new dynamical regimes not observed before, providing deeper insights into the complexity and robustness of GRNs in synthetic biology. These advancements not only enhance our understanding of biological timing mechanisms but also pave the way for innovative applications in the design of synthetic biological systems.

## ACKNOWLEDGMENTS

JS and TA have been supported by grant PID2021-127896OB-I00 funded by MCIN/AEI/10.13039/501100011033 “ERDF A way of making Europe.” JS has been also supported by the Ramón y Cajal grant RYC-2017-22243 funded by MCIN/AEI/10.13039/501100011033 “FSE invests in your future.” RP-C and SM acknowledge funding by the Leverhulme Trust Research Project Grant RPG-2023-085. AH has been supported by the Spanish grant PID2021-125535NB-I00 (MCIU/AEI/FEDER, UE). This research has been also funded through the Severo Ochoa and María de Maeztu Program for Centers and Units of Excellence in R&D CEX2020-001084-M (AH, TA, JS). We thank CERCA Programme/Generalitat de Catalunya for institutional support.

## APPENDIX A: ISOLA FORMATION CRITERION

Consider the case where *q* < *q*_*c*_, where *q*_*c*_ denotes the value of *q* at which the transcritical bifurcation occurs (Fig. 1(b), right panel). For smaller values of *q*, an isola [40] appears. Equivalently, for Eqs. (6), when considering *r* < 0, the isola appears and for *r* = 0, we locally observe a point in the bifurcation diagram, showing its collapse (see Fig. 9). The existence of this isola in the deterministic mushroom bifurcation diagram, together with the existence of the transcritical bifurcation, can be analytically demonstrated through the application of the Morse Lemma [56]:

**FIG. 9.**
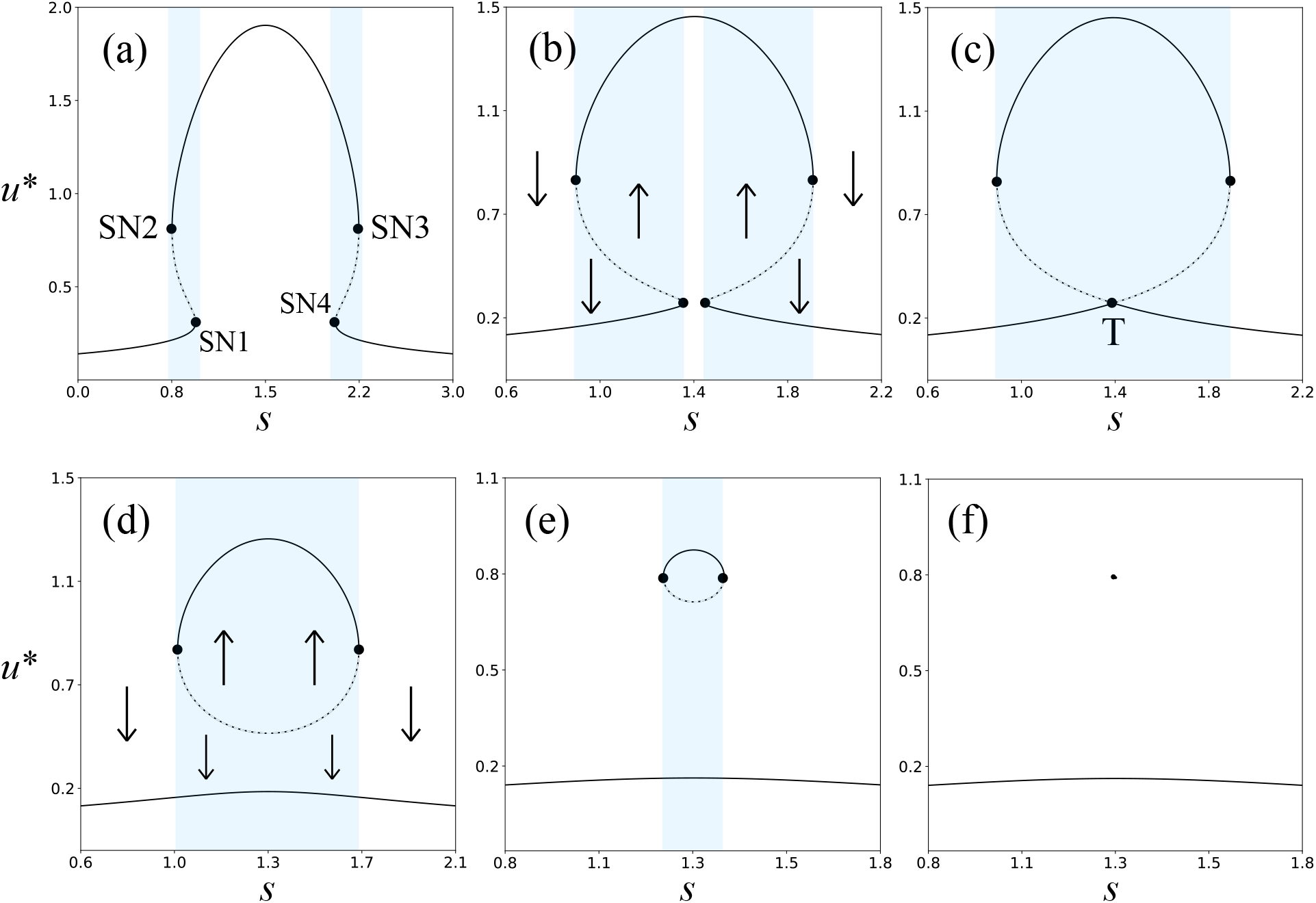
Evolution of the mushroom bifurcation diagram for Eq. (1) at decreasing *q* with *r* = 0.14. Panels (a, *q* = 3) and (b, *q* = 2.786) show the diagram with four SN bifurcations. (c) Transcritical bifurcation at *q* = *q*_*c*_ = 2.784953…. Two examples of isolas are shown with *q* = 2.7 (d) and *q* = 2.605 (e). Panel (f) shows how the isola collapses at *q* = 2.600914. Bistability regions are shown in blue.

### Theorem 1

(**Morse Lemma**). Let *f* : *U* → ℝ be a *C*^3^ func-tion defined on an open set *U* ⊂ ℝ^*n*^. Suppose *x*_0_ ∈ *U* is a non-degenerate critical point (equilibrium point) of *f* with index *s*. Then, there exists a *C*^1^ diffeomorphic immersion *G* : *V* → *U*, where *V* ⊂ ℝ^*n*^ is an open set, such that *g* = *f* ◦ *G* : *V* → ℝ is of the form

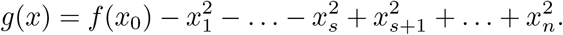

From Morse Lemma, the following corollary can be derived, which provides a criterion for distinguishing between the formation of isolas and transcritical bifurcations based on the sign of the determinant of the Hessian matrix at the critical point.

### Corollary 2.

***(Isola formation and transcritical bifurcation emergence)***. If the determinant of the Hessian matrix of a smooth function *f* (*x, λ*), where *λ* is the parameter at a non-degenerate critical point is positive (negative), then the function exhibits isolas (transcritical bifurcations) in a small neighborhood around the critical point.

*Proof*. By hypothesis, the Hessian matrix *H* of *f* is positive, therefore, *H* is non-degenerate, as it is not zero. By applying the Morse Lemma, we can assert that there exists a local diffeomorphism

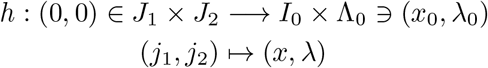

such that *h*(0, 0) = (*x*_0_, *λ*_0_) and 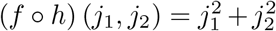, where we assume without loss of generality that ∂_*xx*_*f* > 0, in case it was negative, we define 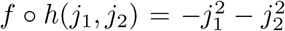. Distinctly as in the case of the transcritical bifurcation, here the level of curve *f* (*x, λ*) = 0 is just a point. The set of points satisfying the equation *f* ◦*h*(*j*_1_, *j*_2_) = 0 forms a set of isolas in the *j*_1_-*j*_2_ plane. Since the transformation *h* is a local diffeomorphism, these isolas map to a set of isolas in the *x*-*λ* plane.

The equation (*f◦h*)(*j*_1_, *j*_2_) = 0 implies 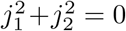, which corresponds to the point (0, 0). Now, if we look at a small neighborhood around (0, 0), the equation (*f ◦ h*)(*j*_1_, *j*_2_) = *c* for small *c* > 0 describes a set of points forming a circle with radius 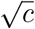 around the origin. This is because for small *c* > 0, 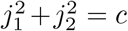 corresponds to a circle with radius 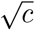 in the *j*_1_-*j*_2_ plane.

When these circles are transformed by *h* into the *x*-*λ* plane, they form isolas in a small neighborhood around the critical point (*x*_0_, *λ*_0_). This is possible because the transformation *h* preserves the topology of the small neighborhood around the origin, and thus the isolas in the *j*_1_-*j*_2_ plane become isolas in the *x*-*λ* plane.

Therefore, if the determinant of the Hessian matrix of a smooth function *f* (*x, λ*) at a non-degenerate critical point is positive, then the function exhibits isolas in a small neighborhood around the critical point. This argument relies on the local structure of the function around the critical point, captured by the Morse Lemma, and the positive determinant of the Hessian matrix ensuring a non-degenerate critical point.

**FIG. B.1.**
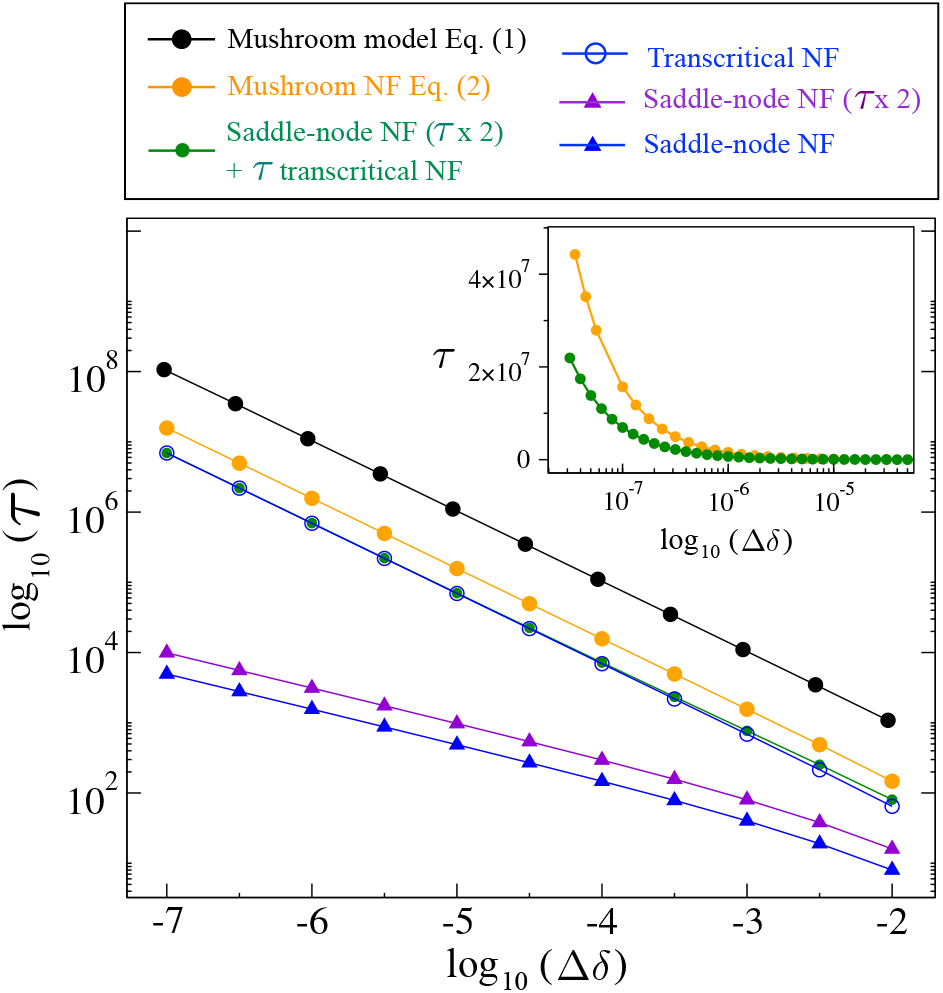
Transient times *τ* close to the bifurcation value (in log-log scale) for six cases given by the: normal form (NF) of the SN bifurcation (blue triangles); twice the times for the NF of the SN (violet triangle); transcritical NF (open blue circles); sum of the times for the transcritical NF plus twice the times of the SN NF (solid green circles); NF of the coupled SN bifurcations collapsing into the transcritical one [Eq. (2)] (orange circles); and the mushroom model given by Eq. (1) (black circles). The inset compares these times (in log-linear scales) for the mushroom NF (orange) and the times assuming additivity of two independent SNs with the transcritical slowing down using their NFs (green).

## APPENDIX B: COMPARISON OF THE TRANSIENT TIMES FOR ISOLATED AND MULTIPLE NEARBY BIFURCATIONS

In this section, we compare the transient times close to several bifurcations by inspecting six different cases. To facilitate this comparison, we define the distance to the bifurcation in a general way as Δ*δ*= |*p*−*p*_*_|, where *p* is the control parameter and *p*_*_ denotes the bifurcation value. The transients have been computed choosing initial conditions passing through the delaying regions for each case [57] e.g., for SN bifurcations we have chosen initial conditions passing through the bottleneck of the ghosts. By doing so, we aim to compare such times to show that the dynamics close to multiple bifurcations, such as the ones studied for the mushroom bifurcations diagrams, are non-additive. As we show below, not only are the effects on transients in the mushroom bifurcation non-additive but also they show a coupling effect of both SN1-SN4 when approaching the transcritical bifurcation. That is, a resonance effect of these bifurcations produces a synergistic slowing down. We first show the results for transients close to a single SN bifurcation using the normal form 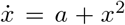 (blue triangles in Fig. B.1) and the transients obtained by the sum of times for two independent and distant SN bifurcations (violet rectangles in Fig. B.1). These later results will be compared with the multiple nearby bifurcations as a case to look for additivity in the resulting transient times. For this case, transients are measured from *x*(0) = −0.1 to *x* = −0.1. Then, the same results are obtained for the normal form of the transcritical bifurcation, 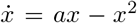 (open blue circles in Fig. B.1). The green data in Fig. B.1, which appear overlapped to the results for the transcritical bifurcation, show the times for the transcritical case plus twice the time of a SN bifurcation. In a linear additive scenario, transient mushroom dynamics phenomena should follow such times. Then, we compute these transients times for the normal form Eq. (2) (orange circles in Fig. B.1 and the gene circuit system given by Eq. (1) (black circles in Fig. B.1). For these two models, we set *s* = 0 and adjusted the opening gap such that Δ*δ* corresponds to the distance 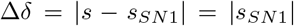. For Eq. (1) we used *r* = 0.14. For the normal form and model of the mushroom bifurcation, transient times were measured for *x*(0) = −0.1 to *x* = 0.1, and from *u*(0) = 0.2 to *u* = 0.4, respectively.

The results show that the longest transients are found for the self-activating gene model Eq. (1) and the normal form (2). These results also show that the slowing down obtained for the multiple bifurcations models departs from an additive effect of individual local bifurcations. For instance, the times obtained with the normal form of the mushroom model and the gene model are much longer than the times obtained by summing up the delays tied to the transcritical bifurcation plus the delays of two independent SN bifurcations. Finally, the times for the multiple bifurcations are about 3-4 orders of magnitude longer than the ones for two single and far away SN bifurcations, either computed separately or additively.

## APPENDIX C: ANALYTICAL STUDY OF THE SCALING LAWS IN THE MUSHROOM BIFURCATION

To analytically study the saddle-node (SN) scaling law for Eq. (1), it is sufficient to investigate general non-degenerate SN bifurcations.

### Theorem 3.

(***Implicit Function Theorem [58]***). Let *f* be a function of class *C*^*p*^ (with *p* > 0) defined on an open set *U* of ℝ^2^ and taking values in R. Let (*x*_0_, *y*_0_) be a point in *U* such that *f* (*x*_0_, *y*_0_) = 0 and such that the partial derivative of *f* with respect to the second variable is non-zero at (*x*_0_, *y*_0_). There exists a real functionφ of class *C*^*p*^, defined on an open real interval *V* containing *x*_0_, and an open neighborhood Ω of (*x*_0_, *y*_0_) in *U* such that, for all (*x, y*) ∈ ℝ^2^:

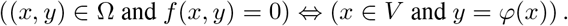

Applying Theorem 3 to the studied model, we can define the following proposition to describe the factorization of the function *f* :

### Proposition 4.

Suppose *f* (*x, r*) is a smooth function with respect to *x* ∈ ℝ in a neighborhood of *x* = 0 such that 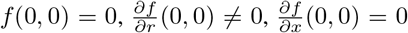, and 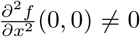. Then, there exists a smooth function *p*(*r, x*) which is nonzero in a neighborhood near the origin *N* = [ −*δ, δ*]^2^, such that *f* can be locally expressed as:

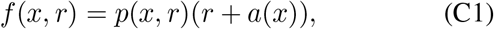

where *a*(*x*) satisfies *a*(0) = *a*′(0) = 0 and *a*″(0)*≠* 0.

*Proof*. Given the condition *f* (*x, r*) = 0 and the orders of vanishing according to each variable for the saddle-node bifurcation through the function *f*, we can apply the Implicit Function Theorem. The saddle-node bifurcation diagram is generated by

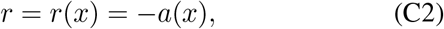

and

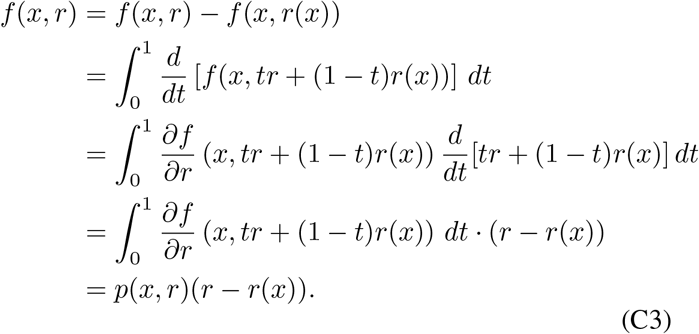

This demonstrates the factorization of *f*. Furthermore, evaluating at the origin and considering that *a*(0) = 0, we get

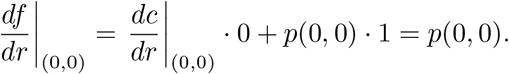

Therefore, at the origin, the derivative of *f* concerning *r* equals the value of the function *p*(*x, r*) at the same point. Moreover, the system’s dynamics in the *r* direction at the origin are governed by the factor *p*(*x, r*), as expected.

Finally, the function *a*(*x*) is a smooth function of degree 2, as derived from Eq. (C2), satisfying *a*(0) = *a*′(0) = 0 and *a*″(0)*≠* 0.

Given these conditions, the scaling behavior near a generic non-degenerate SN bifurcation such as Eq.(C1 with *p*(0, 0) > 0 and *a*″(0) > 0 can be characterized through the following inequality:

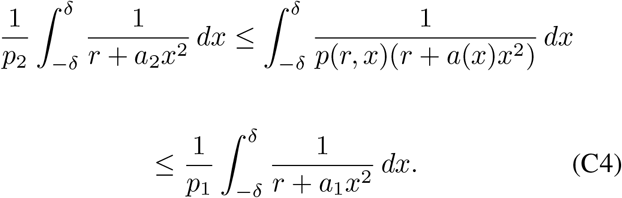

Therefore:

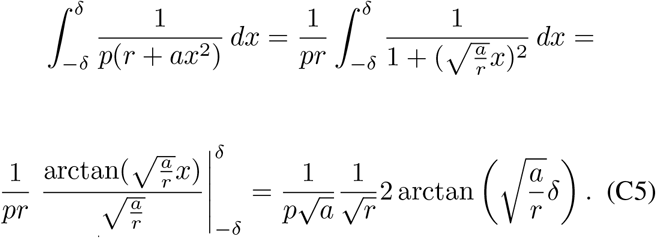

As *r*→ 0^+^, the asymptotic behavior of the system, or equivalently, the scaling law on the transients time near an SN bifurcation is given by

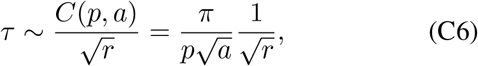

where *C*(*p, a*) denotes a constant pre-factor depending on *p* and *a*. This approximation validates that the integral demonstrates a consistent scaling behavior as *r* → 0^+^, which is independent of the form of the function *p*(*x, r*).

## APPENDIX D: BISTABLE SWITCH MODEL

Similar to the mushroom model given in Section II, we here introduce a bistable switch model to help uncouple the difference between the effects of a single SN bifurcation and multiple.

We use the same simple regulatory model as for the mushroom

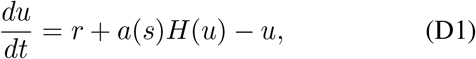

but with a modified external signal *a*(*s*)

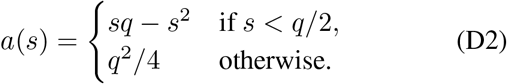

The stochastic models for the bistable switch are the same as for the mushroom bifurcation (see tables I and II) with a change only in transition rate *W*_1_ which is dependent on the new external signal function. In the intrinsic noise case, the signal can be rewritten as

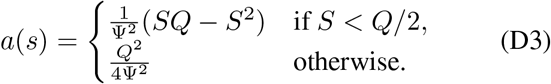

## Notes

### Competing Interest Statement

The authors have declared no competing interest.

